# Collective cell behaviour in mechanosensing of substrate thickness

**DOI:** 10.1101/228478

**Authors:** CG Tusan, YH Man, H Zarkoob, DA Johnson, OG Andriotis, PJ Thurner, S Yang, EA Sander, E Gentleman, BG Sengers, ND Evans

## Abstract

Extracellular matrix stiffness has a profound effect on the behaviour of many cell types. Adherent cells apply contractile forces to the material on which they adhere, and sense the resistance of the material to deformation – its stiffness. This is dependent on both the elastic modulus and the thickness of the material, with the corollary that single cells are able to sense underlying stiff materials through soft hydrogel materials at low (<10 µm) thicknesses. Here, we hypothesised that cohesive colonies of cells exert more force and create more hydrogel deformation than single cells, therefore enabling them to mechanosense more deeply into underlying materials than single cells. To test this, we modulated the thickness of soft (1 kPa) elastic ECM-functionalised polyacrylamide hydrogels adhered to glass substrates and allowed colonies of MG63 cells to form on their surfaces. Cell morphology and deformations of fluorescent fiducial-marker labelled hydrogels were quantified by time-lapse fluorescence microscopy imaging. Single cell spreading increased with respect to decreasing hydrogel thickness, with data fitting to an exponential model with half-maximal response at a thickness of 3.2 μm. By quantifying cell area within colonies of defined area, we similarly found that colony-cell spreading increased with decreasing hydrogel thickness but with a greater halfmaximal response at 54 μm. Depth-sensing was dependent on ROCK-mediated cellular μ contractility. Surface hydrogel deformations were significantly greater on thick hydrogels compared to thin hydrogels. In addition, deformations extended greater distances from the periphery of colonies on thick hydrogels compared to thin hydrogels. Our data suggest that by acting collectively, cells mechanosense rigid materials beneath elastic hydrogels at greater depths than individual cells. This raises the possibility that the collective action of cells in colonies or sheets may allow cells to sense structures of differing material properties at comparatively large distances.

## Introduction

Cells have a complex intracellular apparatus that enables them to mechanosense their material environment. Conceptually, an adherent cell is able to sense stiffness by applying force on a surrounding material and creating a deformation in the material, which is proportional to the material’s intrinsic stiffness. It is thought that cells are able to sense, or to measure, these deformations as a function of the force they exert, and translate them into phenotypic responses. Mechanosensing by cells is now understood to be important or fundamental in a wide range of cellular processes, including division (1, 2), migration (3–5), morphology (6–8), differentiation (9, 10) and mature cell function (11, 12). Despite this, many aspects of the molecular and mechanical control of mechanosensing are not fully understood or are controversial.

The majority of cellular mechanotransduction studies employ polyacrylamide hydrogels functionalised with surface ECM proteins as growth substrates (13). The elastic modulus of this material can be tuned in a convenient manner by adjusting the ratio of monomer to crosslinker (8). This system has been used over a number of years to illustrate cellular phenotypic differences as a function of ECM elastic modulus and is generally understood to be isotropic and to display linear elastic properties (14) (although some experiments suggest in some circumstances its elastic behaviour is nonlinear (15)). Many studies assume that the difference in stiffness experienced by cells is due to the mechanical properties of polyacrylamide, although some groups have suggested that differences in porosity and concurrent variations in the surface binding of ECM proteins may also determine the stiffness of the material that cells detect (16, 17).

Regardless of these questions, it is important to note that the stiffness that a cell measures is dependent not only on the elastic modulus of the material, but also on the material geometry.

This point is best illustrated by phenotypic changes in individual cell behaviour on hydrogels of differing thicknesses adhered to an underlying glass support (18–22). In these studies, single cells remain rounded on soft (eg < 2 kPa) hydrogels above a ‘critical thickness’, but begin to spread progressively more as the hydrogel gets thinner, even though the elastic modulus of the hydrogel is unchanged. An explanation for this behaviour is that the lateral displacement exerted by a cell is constrained by the attachment of the hydrogel to the underlying glass support (23). Most studies find a ‘critical thickness’ (also sometimes called ‘mechanosensing length’) for cells cultured on linear elastic materials of several microns (18, 24–26) but with others predicting longer scales (20, 27). These results are likely to depend on the magnitude of the forces cells exert, the resultant displacements, the size of the cells, and of the dimensions of the focal adhesions (20, 28–30).

In most tissues, however, cells do not exist in isolation but interact with each other intimately, both chemically and mechanically. Specific cell-cell adhesion complexes such as desmosomes and cadherins have evolved in these tissues to mechanically couple the cytoskeletons of adjacent cells. To maintain adherence to the underlying ECM, however, they also must exert tensile force on it to balance the forces they exert on each other. In many cases, the forces that groups of cells transmit to the ECM can become very large, and there is evidence that forces may become integrated over the entirety of a cell monolayer (31). This phenomena was illustrated in experiments by Emerman and Pitelka in the 1970s where epithelial layers growing on floating collagen hydrogels were shown to contract them to ¼ of their starting diameter (32). That groups of cells strain hydrogels to a greater extent than single cells has also been shown by others (33–35). For example, Zarkoob *et al*. found that average hydrogel displacements were five times higher for groups of ~8 cells compared to single cells, with some deformations reaching in excess of 100 µm (33).

The greater lateral displacements that groups of cells impart on ECMs compared to single cells has the implication that cell groups or colonies may be able to ‘feel more deeply’ into matrices than single cells, sensing rigid materials beneath themselves at much greater depths than single cells. This idea is supported by observations that colonies of MDCK cells are insensitive to elastic modulus on polyacrylamide hydrogels of depths < 100 µm (31), but has not to our knowledge been examined quantitatively.

In this study, we explored the idea that colonies of defined sizes sense an underlying rigid support at greater hydrogel thicknesses than single cells. We demonstrate that collective behaviour in cells enables individual cells to interrogate substrate geometry at greater distances than they would be able to do separately, and which suggests that matrix geometry may mechano-regulate behaviour of cell groups.

## Materials and Methods

### Cell culture

Human osteosarcoma (MG63) cells were cultured in Dulbecco’s modified Eagle medium (DMEM) (Lonza, Slough, UK), supplemented with 10% fetal bovine serum (FBS) (Gibco, Life sciences, Paisley, UK) and 1% penicillin-streptomycin (Lonza) in a humidified incubator maintained at 37°C and 95/5% air/CO_2_. MG63 cells were initially plated and passaged in medium using standard tissue culture polystyrene flasks. Media was replaced every 2-3 days, and cells were passaged at 80% confluence. In order to promote single-cell colony formation with sufficient separation between colonies, MG63 were seeded at low density (300 cells/cm^2^).

### Cell characterization

Cell nuclei were visualized by fixing samples in paraformaldehyde 4 % for 20 minutes, before counterstaining with DAPI (4′, 6-diamidino-2-phenylindole) (Fisher Scientific, Loughbourough, UK). Cell proliferation was measured by using the PicoGreen^®^ dsDNA quantitation assay (Fisher Scientific) after 5 days in culture according to the manufacturer’s instructions. Cells were imaged with a Nikon Eclipse Ti inverted microscope (Nikon UK Limited, Surrey, UK). Single cell spreading area was measured in Image J (NIH, Bethesda, MD) by manually drawing an outline around the cell. Colony area was measured in Cell Profiler 2.2.0 (Cambridge, USA) open source software (36). Cell counting was done manually in Image J from the DAPI-stained z-stacks (2 μm steps) / 20× magnification images acquired with the Nikon Eclipse Ti inverted microscope. Z-stacks were analysed using the ‘cell counter’ plugin from the Image J. The number of transient cytoplasm projections was counted manually in Image J for n = 3 different colonies over a period of 3h. In order to measure the thickness of the colonies we performed confocal microscopy (Leica TCS SP8, Cambridge, UK) on DAPI-labelled cells. Colonies (n = 6 for each thin and thick hydrogel) were scanned (2 μm steps, 20× magnification) from the bottom to the top across the thickness of the colony by imaging the DAPI-stained nuclei. Colony thickness was measured using the XZ-scan. By scanning through the entire colony an average intensity profile of the fluorescent signal was recorded. Leica software (LAS X Core Offline version 3.3.0) was used to measure colony thicknesses by analysing the fluorescent intensity profiles. The full-width half-maximum (FWHM) was computed by identifying the two points where the intensity value was greater than 0.2. The thickness of the colony was defined as the distance between these two points in the FWHM of the fluorescent intensity profile at the site of the maximal thickness – thus the values presented indicate the colony maximum thickness.

### Polyacrylamide hydrogel production and characterisation

Polyacrylamide (PA) hydrogels used in this study were prepared using a modified version of the Pelham and Wang protocol (8). In brief, glass coverslips (25 mm diameter, VWR International, Leicestershire, UK), which were used as a rigid support for the PA hydrogels, were cleaned and functionalized with 500 μl, 0.1 M NaOH (Sigma-Aldrich, Gillingham, UK). The coverslips were placed on an 80 °C hotplate for 5 minutes and then washed well with distilled water before drying and adding 500 μl of 100% 3-aminopropyl-triethoxysilane (APES) (Sigma-Aldrich). After another wash, the coverslips were immersed for 30 minutes in 2 ml of 0.5% glutaraldehyde in PBS (Sigma-Aldrich). By varying the concentration ratio between acrylamide (40%) and bis-acrylamide (2%) a range of soft and stiff hydrogels were prepared, for example for 1 kPa hydrogels the concentration of acrylamide was 5% and bis-acrylamide 0.03 %, where for 40 kPa the concentration of acrylamide was 8% and bis-acrylamide 0.48%. We did not directly measure stiffness, but these concentrations of monomer and crosslinker are expected to yield hydrogels of the stated elastic moduli (37).

The mixture was degassed for 15 minutes or longer under a vacuum. In order to initiate polymerization, 1μl of N,N,N’,N’ tetramethylethylenediamine (TEMED; Sigma-Aldrich) and 10 μl of 10% ammonium persulfate (APS; Sigma-Aldrich) were added and the whole solution was vortexed. Thin hydrogels were formed by placing the mixture between a pretreated coverslip with an alkosylated surface (dimethyldichlorosilane, Sigma-Aldrich) and a pretreated glass slide with a hydrophobic surface. Sulfosuccinimidyl 6(4-azido-2-nitrofenyl-amino) hexanoate (sulfo-SANPAH) (Fisher Scientific) and UV light (Chromato-vue TM-20, UVP transilluminator, 240 V) were used to cross-link 500 μl of 0.1 mg/ml fibronectin (Fn) (Merck MiliPore, Watford, UK), to the polyacrylamide hydrogel. The hydrogels were incubated overnight at 4 °C. On the following day, excess solution was aspirated carefully and the PA were washed with PBS.

For thick hydrogels, before the coating step, the hydrogels were washed overnight in excess PBS in a stirred bath order to eliminate acrylamide monomer. To achieve a range of thicknesses, we varied the gelation mixture volume between 6 and 100 μl. Hydrogel thickness was calculated from the difference in the microscope objective z-position for the focal plane of the glass surface contacting the hydrogel and the hydrogel surface; both surfaces being identifiable under phase contrast illumination. PA hydrogels for displacement tracking were also made except that a solution of 10 μm diameter, red fluorescent (580/605 nm) (Fisher Scientific), 2 % (w/v) was also added at a final concentration of 0.002 % (w/v). The microsphere solution was sonicated for 15 minutes prior to addition to the acrylamide/bis-acrylamide mixture.

### Live cell imaging

Time-lapse images were obtained with a Nikon Eclipse Ti inverted microscope (Nikon UK Limited, Surrey, UK) equipped with an automated motorised stage, autofocus and wide-field epifluorescence. Bright field images of cells and fluorescent images of the microspheres were acquired by phase contrast with a 10× plan apo objective. The MG63 were seeded on the Fn-coated PA hydrogels at 300 cells/cm^2^ density, 2 days before the experiment was initiated (to allow formation of small colonies derived from a single cell) inside a humidified incubator maintained at 37 °C and 95/5% air/CO_2_. The microscope chamber was stabilized at 37°C and the atmosphere was enriched with humidified 95/5% air/CO_2_. Time-lapse images were recorded continuously in 18 minutes intervals for 94 h. At each time point bright field and fluorescence (Ex/Em 542/602 nm) images were obtained at one visual field on the PA hydrogel to track cell and hydrogel movement. A total number of 13 colonies cultured on n = 3 hydrogels (thin/thick) were imaged. The colonies were well separated from each other (at least ~900 μm distance one from another) to minimise any displacement interference from μ neighbour colonies.

Pseudopodia, defined here as discrete cellular projections extending outwards >3.5 μm from the edge of a colony were visualised in time lapse movies. The frequency of pseudopodia activity in a colony was determined by counting the number of extensions and retractions associated with single colonies for a time period of 3 hours, commencing 6 h from the start of the time-lapse experiment (n = 5 colonies). Only pseudopodia greater than 3.5 μm were counted. Values are expressed as extensions or retrations/hr.

### Hydrogel displacement tracking

Hydrogel displacements generated by MG63 tractions on PA hydrogels were measured by tracking the displacements of fluorescent microspheres embedded within the hydrogels. These measurements were made using a custom MATLAB (Mathworks Inc., Natick, MA) algorithm based on spatial cross-correlation between fluorescent images pairs (38). This algorithm has been used previously to measure the in-plane hydrogel displacement fields produced by single cells and colonies on PA hydrogels (33). Briefly, a uniform grid of points (56 rows by 48 columns) spaced 3.22 µm (5 pixels) apart was overlaid on the initial image corresponding to t = 0 h. Subwindows (9 pixels x 9 pixels) around each point were then correlated between image pairs in order to track the displacements of the embedded fluorescent microspheres. The grid point positions were then updated and the process was repeated, such that the grid points tracked with the material throughout the course of the experiment. Displacements for a particular timepoint were, unless stated otherwise, measured with respect to t = 0. Thus, displacements were calculated cumulatively.

### Hydrogel displacement evaluation

The mean cumulative magnitude of the displacement for n = 10 colonies on thin and thick hydrogels was determined by calculating the mean of the grid displacements for colonies at a specific time point and subtracting background displacements caused by thermal drift or other sources of noise. Background displacements were determined by averaging the displacement of the top 10 nodes of the grid and the bottom 10 nodes of the grid (nodes unaffected by cell-induced deformations at a region distant from the colony edge). To determine differences in the magnitude of colony hydrogel displacements between thin and thick PA gels, the highest 10 % of the cumulative displacement magnitudes produced by each colony were averaged at each time point of interest.

To assess differences in the extent of hydrogel displacement, average displacements as a function of radial distance from the colony edge were also calculated. This calculation was made by first identifying the colony centre (for an explanation, see Supp. Figure S1A). Next, the radial position of each grid point with respect to the colony centre was calculated. The radial position was divided in equidistant bins from the centre of the colony outwards (Figure S1B) and the cumulative nodal displacements were averaged over all nodes contained within a radial bin identified in the first frame (Figure S1C). Subsequently the average displacement was plotted as a function of the radial distance from the colony edge, defined by calculating the colony radius based on the colony area considered as a perfect circle (Fig. S1D and E).

### Statistical analysis

Statistical significance (p < 0.05) was determined in GraphPad Prism 7 (GraphPad Software, Inc., California, USA) by performing t-tests between two groups with normally distributed data. Data normality was assessed by a Shapiro-Wilk test. For data that was not normally distributed, a Mann-Whitney test for paired data was used. Data are presented as mean ± standard deviation (SD).

## Results

### MG63 cells are mechanosensitive to hydrogel substrate thickness

We first examined and compared the contractile behaviour of individual cells and colonies of the osteosarcoma cell line, MG63, cultured on fibronectin-linked polyacrylamide (PA) hydrogels, ~200 μm in thickness, that varied in elastic modulus (0.5 – 40 kPa). As expected, we found individual MG63 cells were mechanosensitive, with their spread area increasing asymptotically as a function of hydrogel elastic modulus (Supplementary Figure S2). Cells on hydrogels of ~1 kPa bulk modulus did not spread and remained rounded, and so we chose to use hydrogels with this modulus for all further experiments that investigated the effects of hydrogel thickness.

To modulate hydrogel thickness, we simply reduced the acrylamide/bis-acrylamide reaction mixture volume to vary between 10 μ L. This technique produced hydrogels of mean thickness 6.3 μm ± 4 μm for 6 μL hydrogels and 19.5 μm ± 5.0 μm for 10 μm hydrogels, as measured by light microscopy (Supplementary Figure S3). In these thin hydrogels, there was spatial heterogeneity in hydrogel thickness, which allowed us to examine cell spreading across a range of thicknesses.

MG63 cells were plated on these hydrogels and, similarly to a previously published study (18), we found that individual MG63 adopted more spread morphologies on thin compared to thicker hydrogels (Figure 1A) and that cell spread area decreased asymptotically as hydrogel thicknesses increased to ~ 10 μm (Figure 1B). Above this thickness, cell spread area remained constant. The relationship between hydrogel thickness and cell spread area fitted well to an exponential model (*Y = (Y_0_ - Y ∞)e^-kx^ + Y ∞;* R^2^ = 0.960) with a value for ln2/*k* of 3.2 μm (equivalent to half-life in an exponential decay – hereafter referred to as ‘*tactile half-depth*’; Figure 1B) consistent with the ~3.4 μm value reported previously for marrow stromal cells (MSCs) (18).

**Figure 1.**
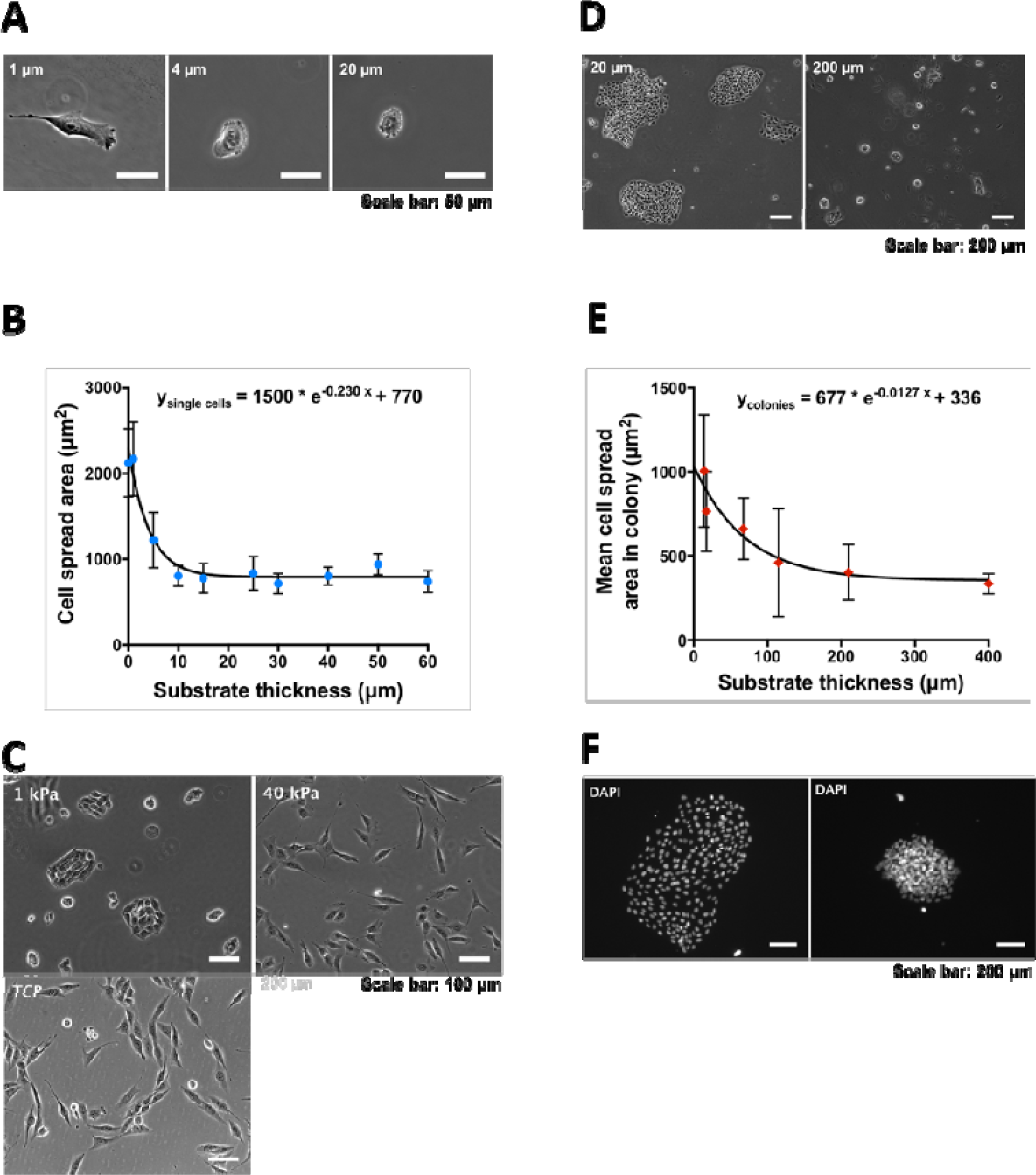
MG63 colonies mechanosense boundaries at greater depths than individual cells. **(A)** On hydrogels with the same elastic modulus (1 kPa), MG63 spread to a greater degree on thin compared to thick hydrogels as shown by phase contrast micrographs. **(B)** Mean cell area decreased asymptotically as thickness increased to a threshold of ~ 10 μm, with the data fitting to an exponential model (R^2^ = 0.960) with a half maximum response (ln2/*k*) at 3 μm. Data presented as mean ± SD for n = 3 – 10 single cells. **(C)** When plated on 1 kPa hydrogels of thickness 200 μm, MG63 cells formed discrete colonies, in contrast to on stiff (40 kPa) hydrogels or tissue culture plastic. **(D)** Colonies cultured on thin hydrogels (~20 μm) appeared well spread, with each cell in the colony visible. In contrast, on thick hydrogels (200 μm) colonies are round and densely packed with cells. **(E)** The area occupied by a cell in a colony decreased asymptotically as a function of thickness. Data presented as mean ± SD for n = 4 – 15 colonies. The data were fitted to an exponential model (R^2^ = 0.916) with a half maximum response of 54.4 μm, and a maximum density at thickness > 200 μm. **(F)** DAPI staining showed that on thin hydrogels nuclei were well separated, but on thick hydrogels nuclei density was greater.

When plated and allowed to proliferate on PA hydrogels with low elastic modulus (< 2 kPa) and a thickness above 20 μm, MG63 cells formed dense, round, discrete colonies after 3-4 days (Figure 1C). In regions of thickness <10 μm on low stiffness hydrogels, cells did not form colonies. On stiffer hydrogels (> 10 kPa), however, MG63 cell morphology was similar to that on tissue culture plastic, with cells appearing mesenchymal in nature, spreading and migrating away from one another, with no colony formation (Figure 1C).

### MG63 cell colonies feel more deeply into hydrogel substrates than single cells

To explore the relationship between colony morphology and hydrogel thickness, we plated MG63 cells at low density on 1 kPa PA hydrogels (300 cells/cm^2^) and allowed individual colonies to form and grow over a period of five days on hydrogels that varied in thickness between ~20 and 400 μm. We found overt differences in the global morphology of colonies relating to hydrogel depth (Figure 1D; and Supplementary Figure S4A). On thin hydrogels colonies appeared well-spread, with each cell in the colony clearly distinguishable. On thicker hydrogels, however, colonies appeared rounded and densely packed. DAPI staining revealed that on thin hydrogels nuclei were well separated, whereas on thick hydrogels the cell nuclei were tightly clustered and difficult to distinguish from each other (Figure 1F). As a result, at a given time point (day 5), colonies on thin hydrogels were significantly greater in total area than colonies on thick hydrogels (6.73 ± 2.4 × 10^5^ μm vs. 4.69 ± 2.1 × 10^4^ μm, respectively at day five, p < 0.0001). A linear relationship between colony cell number and colony area was found on both thin and thick hydrogels. The slope was ~3.5 times higher for thick (3150 [cells/mm^2^]) versus thin (882 [cells/mm^2^]) PA gels (Supplementary Figure S4B) which we interpret as reflecting the more tightly packed nature of the colonies, on thick hydrogels.

As a metric for assessing the relative ‘spreading’ of colonies and the cells within them, we divided the cell number of each colony by the total area of each colony. This gave us the mean area occupied by an individual cell within a colony (equal to the reciprocal of cell density; referred to hereafter as ‘*colony-cell area*’) as a function of hydrogel thickness. For consistency, we defined a colony as occupying an area of between 4 × 10^4^ and 4 × 10^5^ μm^2^. By plotting colony-cell area against hydrogel thickness, we found that this metric decreased as a function of hydrogel thickness (Figure 1E). Again, we fitted the data to an exponential model, and recovered a colony *tactile half-depth* at a thickness of 54.4 μm and a minimum constant colony-cell area for thicknesses > 200 μm (R^2^ = 0.916), an order of magnitude higher than the value obtained for single cells. Together, this data suggests that cell colonies are able to mechanosense boundary effects at greater depths than single cells can.

It is possible that thinner hydrogels may promote proliferation and that this may have contributed to the differences observed, particularly when one considers that stiff hydrogels promote cell proliferation over soft hydrogels (4). In the time period investigated in our experiments, however, no significant differences in proliferation were found between cells on thin or thick hydrogels (4.2 ± 0.8 ng/cm^2^ of DNA compared to 4.6 ± 0.9 ng/cm^2^; p = 0.64, n = 3).

### Thickness mechanosensing in MG63 single cells and colonies is dependent on ROCK activity

Previous studies have indicated that durotactic mechanisms, mediated by cell contractility, may promote cell-cell interactions on soft hydrogel substrates (39, 40). This occurs as a result of neighbouring cells detecting dynamic strains exerted by each other on the hydrogel to which they are attached, and does not depend on direct cell contact. To determine if colony formation is dependent on cell contractility, we examined colony formation in the presence of Y27632, an inhibitor of ROCK and myosin type II contractility. Confirming our hypothesis, colonies did not form on soft hydrogels at any thickness in the presence of the inhibitor, with cells remaining well-separated. This was in contrast to in the absence of the inhibitor, where cells formed tightly packed colonies on soft, thick hydrogels, and more flattened colonies on soft, thin hydrogels (Figure 2).

**Figure 2.**
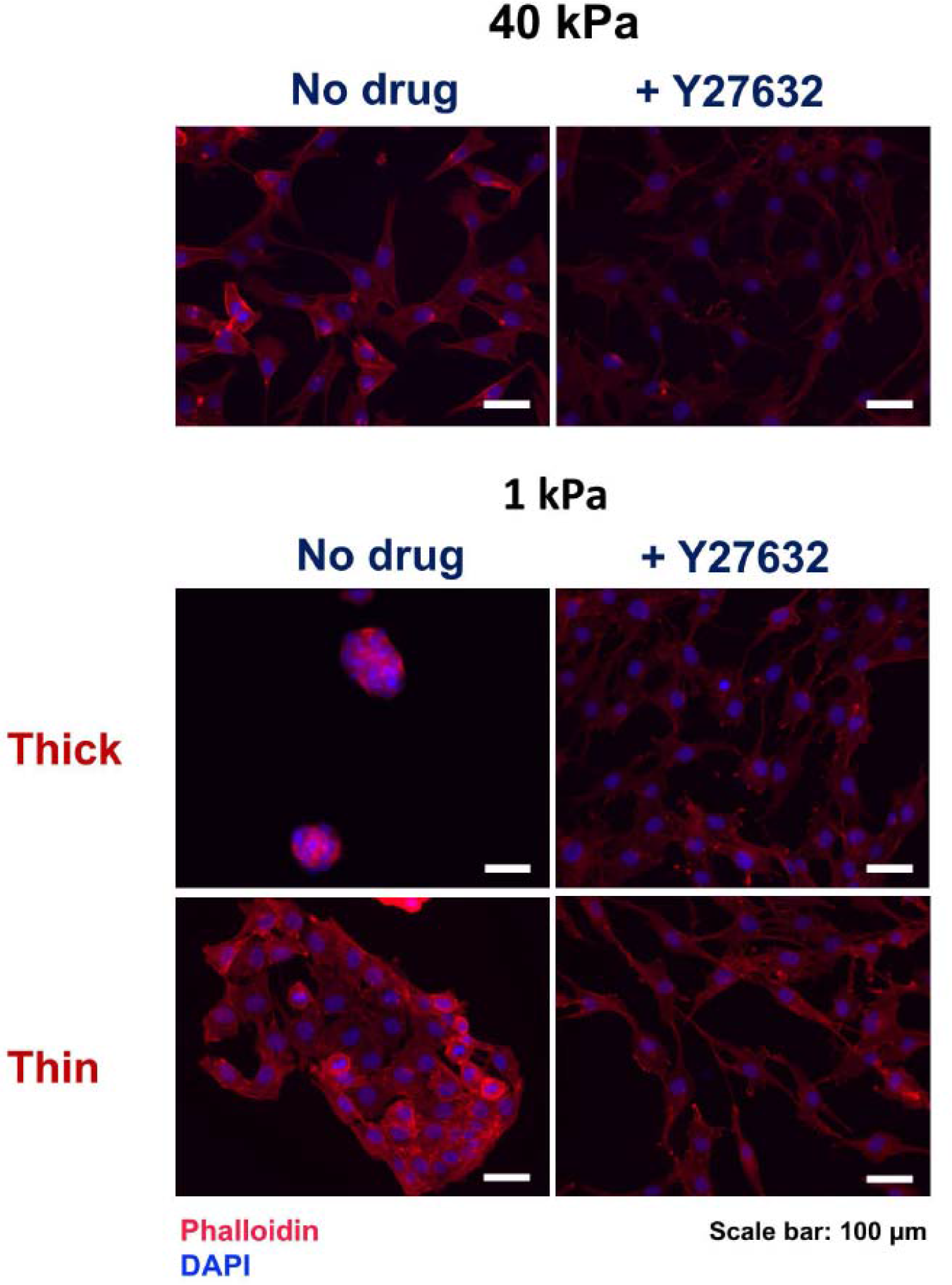
ROCK inhibition abolishes colony formation on soft hydrogels. In the absence of ROCK inhibitor (Y27632) MG63 cells grow as separate, individual cells with a well-defined actin cytoskeleton on stiff (40 kPa) hydrogels. Y27632 addition results in less prominent actin stress fibre formation and more spindle-like cellular projections (compare upper images). On 1 kPa, thick hydrogels MG63 cells grow as rounded, compact colonies, whereas on 1 kPa thin hydrogels, colonies are flattened, with more widely separated cells (phalloidin – red; DAPI – blue; left panels). Y27632 (10 μM) abolishes colony formation on both thin and thick 1 kPa hydrogels with cells appearing similar in morphology to cells on stiff hydrogels (right panels).

### Colony morphology is affected by hydrogel substrate thickness

Phase contrast and epifluorescence imaging of colonies on hydrogels suggested not only quantitative differences in area and cell density, but also qualitative differences in cell organisation as a function of hydrogel thickness. In order to explore this further, we imaged DAPI-labelled cells on ~20 and ~200 μm thickness hydrogels (hereafter referred to as ‘*thin*’ and ‘*thick*’ respectively) using confocal microscopy. On thin hydrogels, smaller colonies (< 4× 10^5^ μm^2^) were dome-shaped, with a flat basal layer and with several cell layers present. μ Larger colonies (> 4.5 × 10^5^ μm^2^) tended to form monolayers and were flatter, consisting of a μ single cell layer (Figure 3A). On thick hydrogels, smaller colonies appeared to form a depression in the hydrogel, which they occupied as a multilayer. This effect was also seen for larger colonies, where a central, densely packed multilayer was always present, with a thinner layer of cells extending beyond the central depression. Mean colony thickness was more than three times greater on thick compared to thin hydrogels (p < 0.05; Figure 3B).

**Figure 3.**
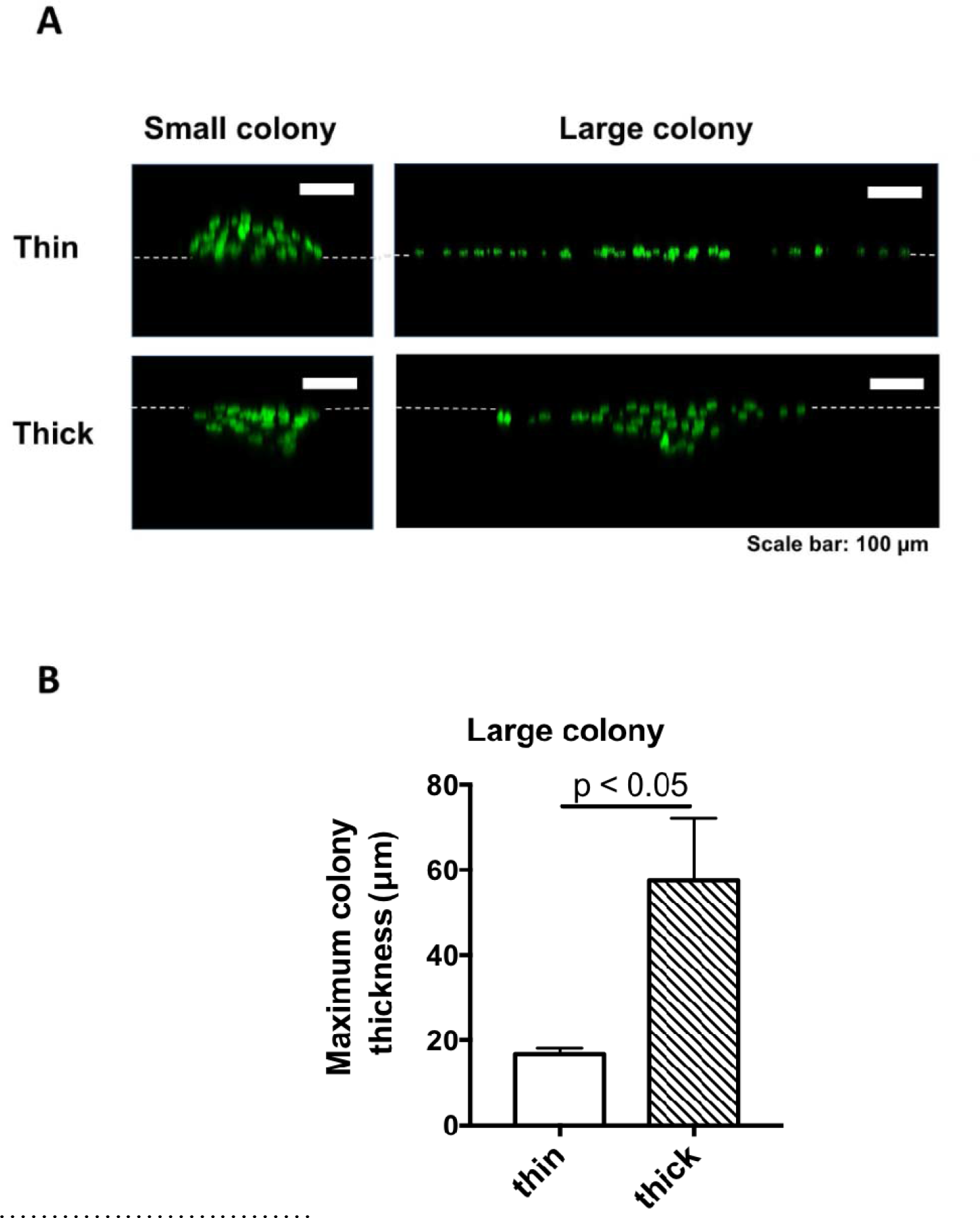
MG63 colonies morphology. **(A)** Small colonies (< 4 × 10^5^ μm^2^) on thin hydrogels were dome-shaped, with a flat basal layer and with several cells layers present, whereas on thick hydrogels colonies appeared to form a depression in the hydrogel, which they occupied as a multilayer. Larger colonies on thin hydrogels (> 4.5 × 10^5^ μm^2^) were flattened, formed of a single cell layer, whereas on thick hydrogels they formed a multilayer depression. These difference in morphology were observed by using laser scanning confocal microscopy on DAPI stained colonies. **(B)** The thickness of the colonies (between 4 × 10^4^ and 4 × 10^5^ μm^2^ in area, n = 4) measured by confocal microscopy was significantly greater on thick compared to thin hydrogels (by a factor of 3.3, p = 0.029). Data presented as mean ± SD of the colony thickness. Statistical significance assessed by an unpaired t-test.

### Colonies on thick hydrogel substrates exert greater contractile displacements than colonies on thin hydrogel substrates

To compare the time course of colony morphology on thin and thick hydrogels, we used time lapse imaging. At 2 days post-plating (0 h), morphology of colonies on both thin (20 μm) and thick (200 μm) hydrogels was similar, but at later time points, colonies on thin materials began to spread to a greater degree than those on thick materials (Supplementary Video 1 and Supplementary Figure S5A and B). In particular, the cells within colonies on thin hydrogels became distinguishable as discrete, phase-dark cells as the colony area increased, indicated call flattening. In contrast, the cells within colonies on thick hydrogels remained indistinguishable in phase-bright areas, indicating compacted/aggregation, even as colony area increased. On thin hydrogels, cells occasionally remained less associated with the colony, but appeared unable to migrate away from the colony, ultimately regaining contact (Supplementary Video 2).

We hypothesised that the differences in colony morphology between thin and thick hydrogels may be a result of differences in the magnitude of hydrogel displacements in response to cell/colony-induced traction forces. In this conceptual model, colonies on both thin and thick hydrogels act to contract the hydrogel (radially displacing the hydrogel surface towards the centre of the colony). However, this contraction is constrained on the thin hydrogels by the proximity of the underlying glass support – a situation which is not true for colonies on thicker hydrogels (13). To test this possibility, we incorporated fiducial fluorescent marker beads (0.5 μm in diameter) in thick and thin hydrogels, and measured colony-induced surface displacements with respect to time. Colony induced displacements in the hydrogels were clearly dependent on thickness (Supplementary Video 3). In general, displacements on thin hydrogels were localised primarily to the regions occupied by cells, whereas on thick hydrogels, displacements extended well beyond the colony periphery (Figure 4A and Supplementary Video 4). On thick hydrogels, displacements were in general directed inward, radially towards the colony centre, while on thin gels displacements were less directional, with both inward and outward displacements (see also Supplementary Video 5, which shows tracking of gel displacements). In addition, the magnitude of the displacements was significantly lower on thin hydrogels compared to thick. For example, after 94 h in culture the mean displacements were 1.9 ± 1.2 μm and 3.9 ± 0.8 μm (p < 0.01) on thin vs. thick hydrogels. This was reflected in a greater frequency of large displacements compared to small displacements for colonies on thick hydrogels vs those on thin hydrogels (Figure 4B). For both thin and thick hydrogels, mean displacement magnitudes increased with respect to time, with significant differences evident from 50 h (Figure 4C). We reasoned that any differences in the displacement may be masked by intrinsic differences in the colony size and cell number between colonies on thin vs. thick over the entire culture period - mean colony area on thin materials being significantly larger at the end of the 94 h analysis period. To correct for this, we next compared displacements around colonies on thin vs. thick hydrogels that did not differ in size significantly (n = 6, p = 0.18) over a ~3 h time period. The magnitude of these displacements was lower on thin hydrogels compared to thick hydrogels for all colony sizes investigated (Figure 4D). We also compared the maximum displacements of colonies on thin vs. thick hydrogels by sampling the highest 10% of displacement values for each frame series and calculating a mean. Over a 94 h imaging period, this metric was significantly lower for thin colonies vs thick colonies (at 94 h, thin: 8.0 ± 3.5 μm, thick 14.8 ± 3.3 μm; for 90 – 94 h p < 0.001; for 8 – 90 h p < 0.01; for 2 - 8 h p < 0.05 and for 0 - 2 h p = 0.105; Supplementary Figure S5C).

**Figure 4.**
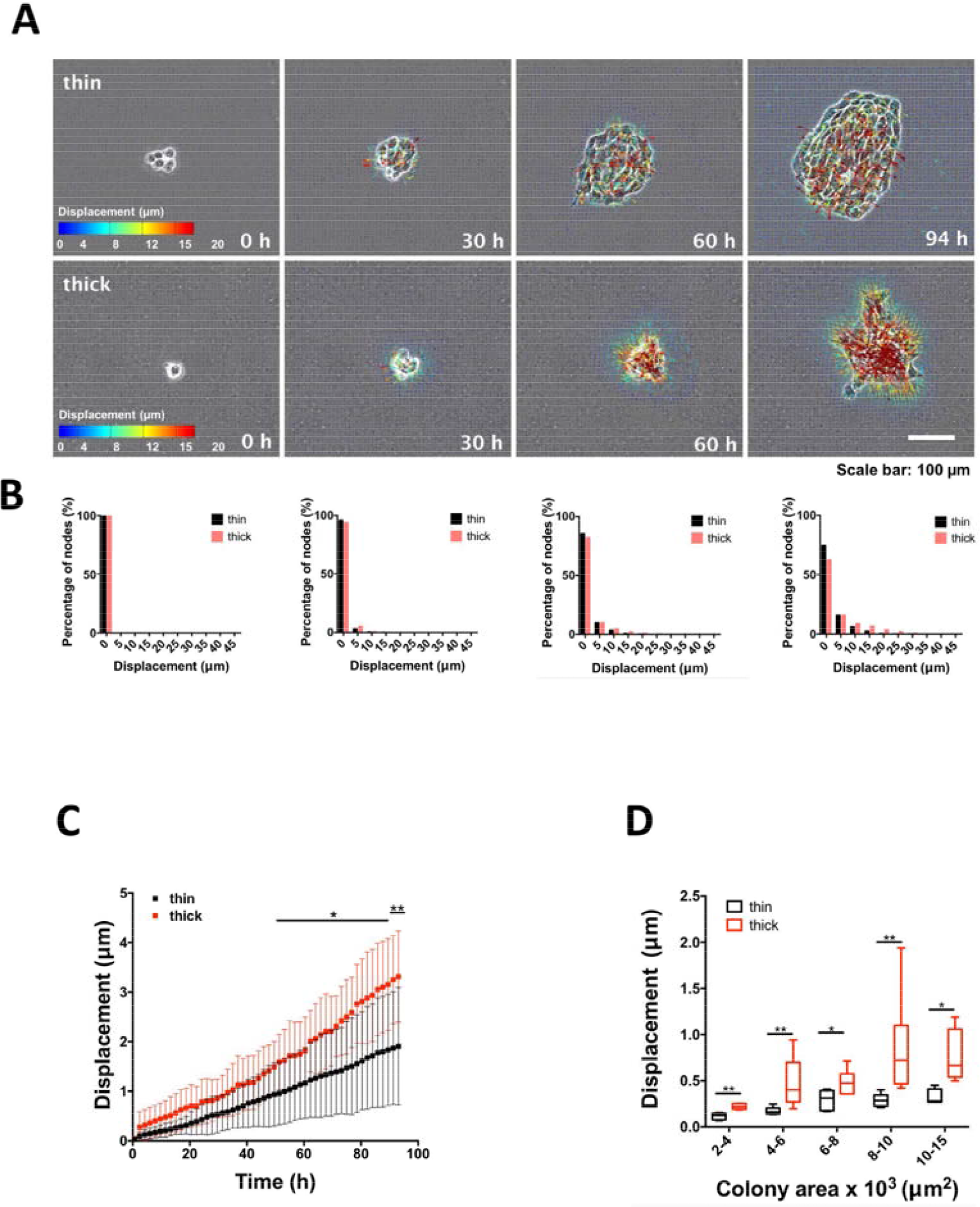
Displacements during MG63 colony formation on 1 kPa fibronectin-coated PA hydrogels. **(A)** In colonies on thin hydrogels, displacements (vectors and their magnitude indicated by coloured arrows) were localised primarily to the regions occupied by cells, whereas in colonies on thick hydrogels, displacements extended greater distances from the colony edge (see also Suppl. Video 4). **(B)** Displacements of larger magnitude were more frequent on thick compared to thin hydrogels, as illustrated by histograms showing the displacement frequency of a given magnitude. **(C)** Mean hydrogel displacements increased with time and were greater in magnitude on thick compared to thin hydrogels (n=10, significant differences in mean displacement occurred after 50 h in culture at 94 h, thin: 1.9 ± 1.2 μm, thick 3.9 ± 0.8 μm, **p < 0.01 for 90 – 94 h, *p < 0.05 for 50 – 90 h). Data presented as mean ± SD of the colony displacement. Statistical significance assessed by Mann-Whitney U test. **(D)** When comparing colonies of equal area, displacements over a period of 3 h were significantly greater on thick hydrogels compared with the thin hydrogels. Data presented as mean ± SD of the colony displacement, n = 5. Statistical significance assessed by Mann-Whitney U test.

### Displacements extend greater distances from the periphery of colonies on thick hydrogel substrates compared to those on thin hydrogel substrates

In addition, mean displacements were not only greater on thick vs. thin hydrogels, but also extended a greater distance from the colony periphery, as shown in displacement contour maps (Figure 5A). By calculating the magnitude of the displacements at increasing distances from the colony edge, we found that displacements declined with respect to distance from colony edge. The decays were fitted to an exponential function, with a half-maximal (ln2/*k*) response of 10.3 μm for thin compared to 25.7 μm for thick hydrogels (Figure 5B).

**Figure 5.**
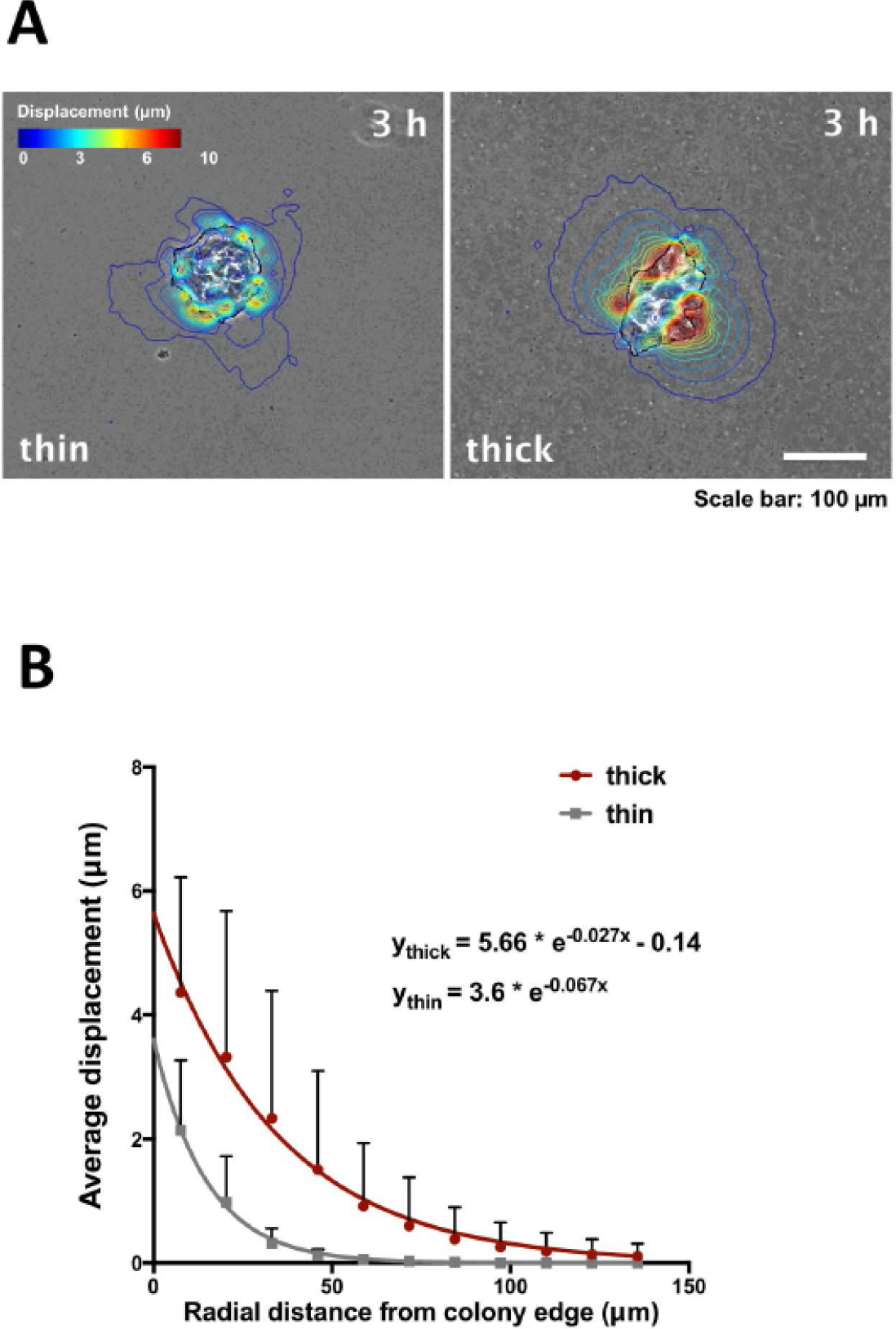
Displacements for colonies on thick hydrogels extended further from the colony periphery than those on thin hydrogels. **(A)** Contour plots of the hydrogel displacement after 3 h for same size colonies. Regions of high displacement are localised at the edge of the colony on thick hydrogels; the magnitude of the displacement is greater on thick than on thin hydrogels. **(B)** Displacements propagated further from the colony edge on thick hydrogels than on thin hydrogels. Data were fitted to an exponential model (R^2^ _thin_ = 0.995, R^2^ _thick_ = 0.998) Data presented as mean ± SD, n = 6.

### Extension and retraction of pseudopodia is more frequent in colonies on thick compared to thin hydrogels

We also observed other differences in colony behaviour by time-lapse microscopy. First, the fiduciary beads beneath a colony moved out of the focal plane of the hydrogel surface during an imaging experiment, reflecting our earlier confocal data and indicating that colonies on thicker hydrogels form depressions (this was not observed on thin hydrogels; Supplementary Video 6). On colonies on thick hydrogels, we observed the frequent extension and retraction of cytoplasmic/cellular extensions, or pseudopodia, at the colony periphery, which was less obvious in colonies on thin hydrogels (Figure 6A). In parallel, we observed deformations consistent with a ‘pinching’ of the hydrogel; this was evident visually on fiduciary bead images as an alignment of marker beads in a direction radial to the colony centre (Figure 6A). In the vicinity of these ‘pinches’ there were large, local displacements occurring orthogonally and directed towards the pinch (Figure 6B; Supplementary Video 6). The frequency of extension and retraction of these pseudopodia was significantly (p = 0.026) greater for colonies on thick hydrogels vs those on thin hydrogels (Figure 6C). Although this phenomenon was associated with pseudopodium extension (83 % of ‘pinches’ corresponded to pseudopodia), they were not observed to extend over the full length of the ‘pinch’, with the deformation extending a mean distance of 59 ± 14 μm pseudopodium. We cannot however exclude that our calculations are limited by the resolution of phase contrast microscopy, and that, thin, microscopically invisible pseudopodia do not extend the full length of a ‘pinch’. Finally, pseudopodia appeared to act to connect adjacent colonies and act to promote the coalescence of colonies on thick hydrogels (Figure 6D and Supplementary Video 7). When colonies were in close proximity, pseudopodia were often observed transiently extending towards a neighbouring colony. This resulted in the connection of cellular bridges between colonies, which subsequently promoted the coalescence of the colonies.

**Figure 6.**
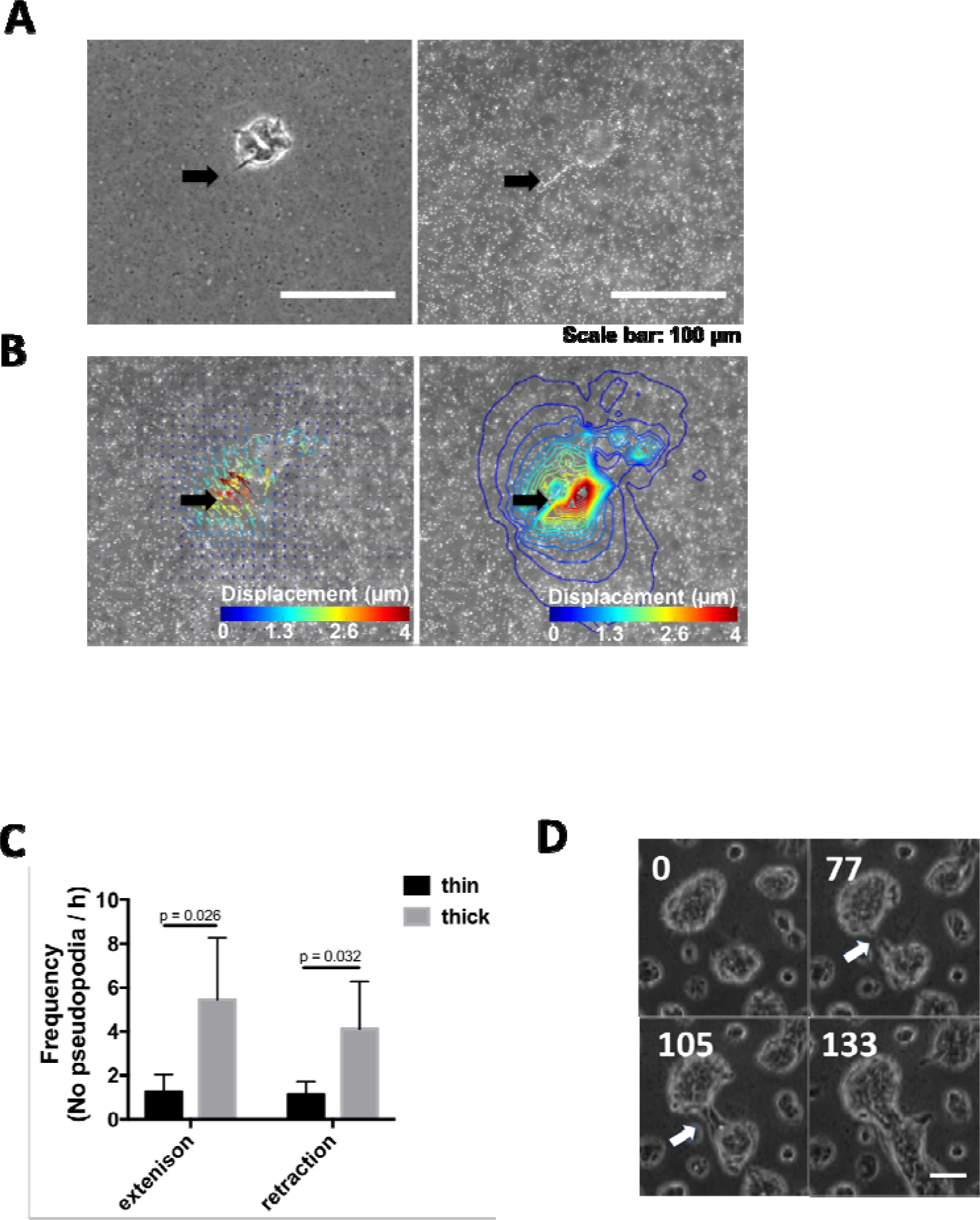
Pseudopodia are responsible for localised, transient and extensive deformations. **(A)** On thick hydrogels, frequent extensions and retractions of cytoplasmic extensions, or pseudopodia, at the colony periphery were observed on the phase contrast images (see also Suppl. Video 6). This was less obvious on thin hydrogels. **(B)** The red arrows and the contour plots show that large displacements are localized near the pseudopodia. (**C**) The number of cytoplasmic extensions and retractions observed over a period of 3 h was significantly greater (p = 0.026, p = 0.032) for colonies on thick hydrogels vs those on thin hydrogels. Data presented as mean ± SD, n = 5 colonies. Statistical significance assessed by unpaired t test. **(D)** In neighbouring colonies, pseudopodia occasionally permitted contact between colonies, which acted to promote colony coalescence (see also Supp. Video 7).

## Discussion

In this study, we have shown that cells act as collective groups to detect rigid materials beneath elastic hydrogels. This enables cells that inhabit colonies to detect boundaries at greater distances than they would if acting alone as single cells. This integrative phenomenon may enable collective groups of cells to probe their environment mechanically in a range of biological contexts, including wound healing, cancer metastasis, patterning in development and vasculogenesis.

We first observed that MG63 cells formed discrete colonies on soft hydrogels above ~ 20 μm in thickness. This was in contrast to on tissue culture plastic or on stiff hydrogels, where cells adhered, divided and migrated, leading to an even distribution of separated cells on the growth substratum. Guo *et al.* observed a similar phenomenon, where 3T3 fibroblasts preferentially formed cell aggregates on soft hydrogels (39). This could be explained simply as a surface-tension-like effect, where cells adhere to each other more tightly than to the underlying matrix and aggregate as a consequence. However, as inhibition of cell contractility via ROCK abolished colony formation, like Guo *et al*, we consider it likely to depend additionally on durotactic mechanisms; *i.e.* preferential migration of non-contacting cells towards each other due to the dynamically stretched substratum (3, 39, 40). This mechanism has been shown previously to promote cell-cell interactions on soft PA hydrogels (40). In our study, we observed similar phenomena between colonies - nearby but separated colonies on soft substrates coalesced, and cells which became detached from colonies appeared to be prevented from subsequently migrating away from their ‘parent’ colony. These observations all support the notion that cells and colonies communicate mechanically through the compliant hydrogel. Further to this, we found that colony formation was sensitive to hydrogel thickness - MG63 cells did not form colonies on thin regions (< 10 μm) of soft hydrogels. This cut-off point was similar to the hydrogel thickness below which we found cells begin to detect the underlying stiff glass support (‘tactile half-depth’), and similar to those reported in similar studies (18, 20, 29). Together, these data indicate that cells are able to detect dynamic (cells) and static (underlying) objects through compliant hydrogel matrices by a process of active mechanosensing.

Colonies that formed on both 20 μm and 200 μm hydrogels were initially morphologically similar, but as they grew larger, their morphologies began to diverge, with the former spreading to a greater extent than the latter. We attempted to measure differences in cell colony ‘spreading’ by a method analogous to that for single cells. As a general, unbiased metric we simply divided the area of each separate colony by the number of cells (nuclei) within each colony. In doing so, we found a similar relationship between colony ‘spreading’ and hydrogel thickness to that observed in single cells, but with a much greater ‘tactile half-depth’ for colonies in comparison to that for single cells. We concurrently found that that surface displacements in the region of cell colonies on soft hydrogels are much greater than for individual cells, as previously shown in keratinocytes (33).

This data suggests that how deeply a colony ‘feels’ (colony ‘tactile half-depth’) is likely to be related to the magnitude of surface displacements as a function of colony-exerted traction force (essentially the stiffness of the hydrogel that the colony ‘measures’). For larger traction forces the strain field will extend deeper within the gel, thus surface displacements will be affected by the constraints posed by the fixed substrate at greater depth (25), which will result in larger cell-or colony stiffness ‘measurements’. As demonstrated by Sen et al (25) this also means that depth sensing will be reduced as stiffness increases as cells or colonies do not deform stiff hydrogels as much as soft hydrogels. This data is supported by observations that large colonies several millimetres in diameter grown on hydrogels ~ 100 μm in thickness are insensitive to substrate stiffness (41). Our data also support theoretical predictions that cell colonies or sheets might be considered to some extent as contractile units, and that depth sensing is determined by the lateral dimension of the cell sheet in relation to the substrate thickness (30), an idea that has been explored experimentally for single cells (20). It will be interesting in future studies to explore whether the elegant and simple ‘line-tension’ model, established by Oakes *et al*. (42) and which can predict traction force localisation for single cells, scales for cell colonies, or whether local actions of groups of cells within a colony or sheet or individual cells are responsible for mechanosensitivity. Although we did not test it, it is also likely that in our system, displacements and traction forces scale with colony size, as shown by Mertz *et al.* (34) and Trepat *et al.* (31). From this it follows that the ‘tactile half-depth’ of colonies may also scale in relation to their size, which could be tested by, for example, controlling colony size more precisely. This might be achieved by microcontact printing matrix proteins (43), or by physically only allowing cell attachment in discrete areas by using stencils (44). These techniques may also uncover information on the dynamics of colony spreading. It is known that single cells reach a steady-state spread area 2-3 hours after plating (45), and that the rate at which this occurs is affected by stiffness. It will be interesting in future studies to investigate this for colonies, by this control of colony size, concurrent with inhibition of cell growth.

As a caveat to the prediction that tactile half-depth might scale with colony area, it is important to note that colonies differed in morphology qualitatively as they increased in size. The method we used to determine ‘colony spreading’ is imperfect and overlooks these qualitative differences. For example, it ignores the fact that cells may not be in contact with the underlying matrix – cells formed layers several layers thick in colonies on thick, soft hydrogels – and does not take into account differences in colony shape. As suggested in another recent study (35), we observed that colonies of cells on soft hydrogels formed cup-shaped pits, or depressions, in the underlying hydrogel. By using 3D traction force microscopy (TFM), this group confirmed that this was due to a cell cluster contracting the hydrogel upwards and centripetally at the cluster periphery, while pushing the underlying gel downwards beneath it – a phenomenon which induced a traction moment in the hydrogel, and which has been observed at a smaller scale for individual cells (46). We were unable to use 3D TFM in our experiments, which would be a prerequisite for computing traction forces around colonies. In our epifluorescence images, displacements in the *z*-plane led to the movement of the hydrogel fiducial markers beyond the depth of field of the 10× objective lens used for acquiring data (~8 μm). Due to this kind of deformation, the in-plane displacements we report for thick gels are likely underestimations of the real, 3D hydrogel deformations induced by the colonies. Note that this is not true, however, for thin or stiff gels, where we did not detect distortions of the gel in the *z*-plane. Despite these limitations, the conclusion we have drawn are not altered, as the displacements are, if anything, underestimated by these methods. Future studies may employ other methods of computing traction forces in three dimensions following accurate marker displacement measurements using confocal microscopy, or using other computation techniques including finite element modelling (35, 47, 48).

Surface displacements decayed exponentially as a function of distance from the colony edge, with displacements extending further from the periphery of colonies on thick hydrogels compared to those on thin hydrogels, despite equivalent colony area. Using finite element simulations Sen *et al.* (25) also predicted exponential displacement decays, with a half-maximal response at ~ 5 μm for contractile ‘stem cells’ of *r* = 40 μm. For cell colonies, we found much large half-maximal responses at 23 μm, supporting the notion that groups of cells exert a greater contractile response than individual cells, a finding that is also supported experimentally by Zarkoob et al (33). Our data again underscores that the extent of displacements is constrained in thin gels by the attachment to the underlying rigid glass support.

We also observed prominent ‘pseudopodia’ transiently extending outwards from the periphery of cell colonies, with significantly more extensions on thick hydrogels compared to thin hydrogels. In the vicinity of these pseudopodia, there were rapid, large displacements in the hydrogel, leading to what appeared to be a ‘pinching’ effect in the hydrogel. Pseudopodia have been shown to exert comparatively large traction forces (47, 49), and so this phenomenon may allow individual cells to ‘sample’ local spatial differences in hydrogel stiffness. We were not able to determine whether these deformations resulted from a pseudopod contracting the gel centripetally, or from a downward motion – high resolution confocal imaging would be necessary to answer this question. We suspect that due to the large size of the pseudopodia, they may comprise a significant portion of an individual cell’s body. In adjacent colonies, the ‘reaching’ behaviour of these pseudopodia like extensions appeared to act to connect and draw neighbouring colonies together, and it is tempting to speculate that a peripheral ‘reaching’ colony cell may be able to exert greater displacements on the underlying matrix than an isolated cell by virtue of the strong contractile forces it exerts being balanced by the colony to which it remains adherent. Future high-resolution time-lapse imaging would be necessary to confirm this.

Our study has several limitations. We have only examined the response of a single transformed cell line, and future studies will be necessary to determine whether the effects we measured are specific to this cell type or are characteristic of a more global phenomenon. Related observations on 3T3 cells (39), primary human keratinocytes (33) and human continuous epithelial cells (35) which, although not designed to determine cell responses to hydrogel geometry, suggest common characteristics of cells on soft matrices, however. Of particular interest will be the comparison of cell types which comprise epithelia (e.g. skin keratinocytes, gut epithelium) with those that do not (marrow stromal cells, skin fibroblasts), and transformed vs. normal cells.

All of our studies were conducted on PA hydrogels functionalised by fibronectin. The majority of published studies agree that PA displays linear elastic properties, making it an excellent material for controlling and measuring the mechanical microenvironment around cells. *In vivo*, of course, cells inhabit three-dimensional materials with nonlinear properties, such as collagens and laminins, so it should always be kept in mind that PA can only represent an idealised, *in vitro* system for deconstructing cellular mechanosensing under well-understood conditions, rather than providing faithful representations of the *in vivo* situation. ECM materials, such as collagen, are known to enable long-range mechanocommunication between cells, artificial tissues and objects or underlying rigid substrata (22, 50–53) and the concept of mechanical communication between tissues through ECMs has been known about for more than sixty years (54). Despite this awareness, the phenomenon remains poorly understood, even though it may play a fundamental role for cells and tissues in gaining positional information in many contexts including patterning in development, wound healing and tissue repair and metastasis. Our study now provides a framework for testing how spatial differences in geometry affect the collective mechanosensing behaviour in cell layers.

## Author contributions

NDE, CGT, BGS, EG, SY designed the study, CGT, YHM, DAJ performed the research. All authors analysed the data, NDE, CGT, BGS and EG wrote the paper.

## Acknowledgements

NDE and CGT acknowledge funding from Wessex Medical Research and the Rosetrees Trust. EG acknowledges a Research Career Development Fellowship from the Wellcome Trust. EAS acknowledges support from the National Science Foundation (CAREER 1452728). We also acknowledge Ben Fabry for helpful discussions.

